# *Baldspot/ELOVL6* is a conserved modifier of disease and the ER stress response *ELOVL6* modifies ER stress

**DOI:** 10.1101/261206

**Authors:** Rebecca A. S. Palu, Clement Y. Chow

## Abstract

Endoplasmic reticulum (ER) stress is an important modifier of human disease. Genetic variation in genes involved in the ER stress response has been linked to inter-individual differences in this response. However, the mechanisms and pathways by which genetic modifiers are acting on the ER stress response remain unclear. In this study, we characterize the role of the long chain fatty acid elongase Baldspot (ELOVL6) in modifying the ER stress response and disease. We demonstrate that loss of *Baldspot* rescues degeneration and reduces IRE1 and PERK signaling and cell death in a *Drosophila* model of retinitis pigmentosa and ER stress (*Rh1^G69D^*). Dietary supplementation of stearate bypasses the need for Baldspot activity. Finally, we demonstrate that *Baldspot* regulates the ER stress response across different tissues and induction methods. Our findings suggest that *ELOVL6* is a promising target in the treatment of not only retinitis pigmentosa, but a number of different ER stress-related disorders.

**AUTHOR SUMMARY:** Differences in genetic background drives disease variability, even among individuals with identical, causative mutations. Identifying and understanding how genetic variation impacts disease expression could improve diagnosis and treatment of patients. Previous work has linked the endoplasmic reticulum (ER) stress response pathway to disease variability. When misfolded proteins accumulate in the ER, the ER stress response returns the cell to its normal state. Chronic ER stress leads to massive amounts of cell death and tissue degeneration. Limiting tissue loss by regulating the ER stress response has been a major focus of therapeutic development. In this study, we characterize a novel regulator of the ER stress response, the long chain fatty acid elongase Baldspot/ELOVL6. In the absence of this enzyme, cells undergoing ER stress display reduced cell death, and degeneration in a *Drosophila* disease model. Feeding of excess fatty acids increases degeneration to original disease levels, linking the regulatory activity of Baldspot to its enzymatic activity. Finally, we demonstrate that Baldspot can alter the ER stress response under a variety of other ER stress conditions. Our studies demonstrate that Baldspot/ELOVL6 is a ubiquitous regulator of the ER stress response and is a good candidate therapeutic target.

## INTRODUCTION

Phenotypic heterogeneity is common in simple and complex diseases and is the driving force behind the Precision Medicine Initiative [1-4]. Genetic variation among individuals accounts for much of this heterogeneity, but the identity and nature of modifying genes or variants is largely unknown [4, 5]. These modifying variants are often cryptic and may not influence the physiology or visible phenotypes of healthy individuals, but can alter the expression of disease phenotypes. Understanding the role of modifiers and the pathways in which they function will enable the development of patient-specific therapeutic approaches.

One such process is the response to endoplasmic reticulum (ER) stress, which occurs when misfolded proteins accumulate in the lumen of the ER. This activates the Unfolded Protein Response (UPR) in an attempt to return the cell to homeostasis. Failure to do so will eventually result in apoptosis [6]. The UPR is controlled by the activation of three sensors located in the ER membrane: IRE1, PERK, and ATF6 [7]. IRE1 is the most highly conserved of these sensors and contains an endonuclease domain responsible for the non-canonical splicing of *Xbp1*. The spliced *Xbp1* transcript is translated and the protein translocates to the nucleus, where it activates the expression of chaperones and other genes involved in resolving ER stress [8, 9]. IRE1 also cleaves a number of additional mRNA targets, which reduces the quantity of newly translated proteins into the ER. This process is known as Regulated IRE1 Dependent mRNA Decay (RIDD) and is commonly linked with increased cell death [7, 10, 11]. When PERK is activated, it phosphorylates the translation initiation factor eIF2α, which decreases the translation of most mRNA transcripts with canonical translation initiation mechanisms [7]. This reduces the protein-folding load of the ER and allows for the upregulation of select transcripts involved specifically in the UPR such as ATF4 [7]. ATF6, the third transmembrane activator of the UPR, is transported to the Golgi upon the sensing of misfolded proteins and cleaved, whereupon the cytosolic portion of ATF6 travels to the nucleus to act as a transcription factor, binding ER stress response elements to further upregulate key players in the UPR [7].

Chronic ER stress leads to apoptosis, tissue degeneration, and dysfunction. In rare cases, primary mutations of key ER stress pathway components cause Mendelian syndromes [12-16]. More commonly, ER stress can exacerbate disease, including obesity [17], neurological disease [18], retinal degeneration [7], and some cancers [19]. Experimental genetic or pharmacological manipulation of ER stress levels can alter disease phenotypes, suggesting that inter-individual differences in ER stress responses may influence disease severity. Indeed, human [20], mouse [21], and *Drosophila* [22, 23] show extensive genetic variation in their response to ER stress. Understanding the role of genetic variation in modulating ER stress pathways may provide more therapeutic targets and improve the accurate identification of high risk patients.

In a previous study, we crossed the *Rh1^G69D^* model of retinitis pigmentosa (RP) into the *Drosophila* Genetic Reference Panel (DGRP) to identify genetic modifiers of this disease [23]. In this model, retinal degeneration is induced by overexpression of misfolded rhodopsin in the developing larval eye disc [24], resulting in chronic ER stress and apoptosis. The DGRP is used to study the genetic architecture underlying complex traits. There are approximately 200 inbred DGRP strains, each strain representing a single, wild-derived genome. Thus, the DGRP captures genetic variation that is present in a natural, wild population. Importantly, we have the whole genome sequence of each strain, allowing phenotype-genotype studies [25]. The degree of degeneration induced by this model varied substantially across the strains of the DGRP. Using an association analysis, we identified a number of promising candidates that modify this RP phenotype and have putative roles in the ER stress response [23].

In this study, we demonstrate that *Drosophila Baldspot*, one of the candidate RP modifiers identified in our genetic variation screen, is both a disease modifier of ER stress-induced retinal degeneration and a more general modifier of the ER stress response across different physiological contexts. *Baldspot*, which is orthologous to mammalian *ELOVL6*, is an ER-associated long chain fatty acid elongase which catalyzes the extension of palmitate to stearate [26, 27]. We show that loss of *Baldspot* rescues retinal degeneration by reducing ER stress signaling. Loss of Baldspot elongase activity reduces IRE1 and PERK activity and downstream apoptosis, without affecting the misfolded protein load. Strikingly, in the absence of Baldspot activity, dietary supplementation of stearate bypasses the need for Baldspot to increase UPR activation. Finally, we show that Baldspot regulates the ER stress response in a number of different genetic and chemically-induced ER stress models across multiple tissue types, making *Baldspot* a potentially wide-reaching modifier of ER stress-associated diseases. Understanding how modifier genes alter disease outcomes is the first step to developing personalized therapies.

## RESULTS

### A SNP in *Baldspot* is associated with *Rh1^G69D^*-induced retinal degeneration

In previous work, we overexpressed a misfolded rhodopsin protein in the developing eye disc (*GMR-GAL4>UAS-Rh1^G69D^*), the larval precursor to the *Drosophila* retina and eye, in the 200 strains from the DGRP [23]. Using a genome-wide association approach, we identified SNPs that are associated with degree of retinal degeneration. One of these candidate SNPs, Chr3L:16644000 (BDGP R5/dm3), is located in intron four of *Baldspot. Baldspot* is the only *Drosophila* orthologue of mammalian *ELOVL6*, a member of the very long chain fatty acid elongase family of enzymes (Fig 1A) [28]. Strains carrying the minor allele (C, 18880 ± 3010 pixels) exhibit more severe *Rh1^G69D^–*induced retinal degeneration as compared to strains carrying the major allele (T, 21822 ± 2294 pixels; P < 0.005) (Fig 1B). This is also apparent when examining the qualitative range of degeneration in strains containing each of these alleles (Fig 1C). Based on the position of this SNP in an intron, we predict that it will affect the timing or levels of *Baldspot* expression. We also predict that it is exerting its effect in the small subset of cells that are *GMR*-expressing. This is very difficult to assess, as these cells make up a very small proportion of the eye disc. It is also difficult to know when, during development, this SNP is affecting expression. Nevertheless, we examined the effect of this SNP on *Baldspot* expression in adult flies and in the larval eye/brain-imaginal disc complex. Expression of *Baldspot* is unaffected by variation at this SNP in whole adult flies (not containing the *Rh1^G69D^* transgene) (Fig S1A). In larval brain-imaginal disc complexes, there is not a significant increase in *Baldspot* expression from strains carrying the C allele and expressing the *Rh1^G69D^* transgene (C, 1.15 ± 0.03 as compared to T, 1.01 ± 0.09;) (Fig S1B). While there is not a substantial difference in expression between the alleles, the possibility remains that altering the expression or activity of *Baldspot* may impact degeneration and the causative pathways.

**Fig 1.**
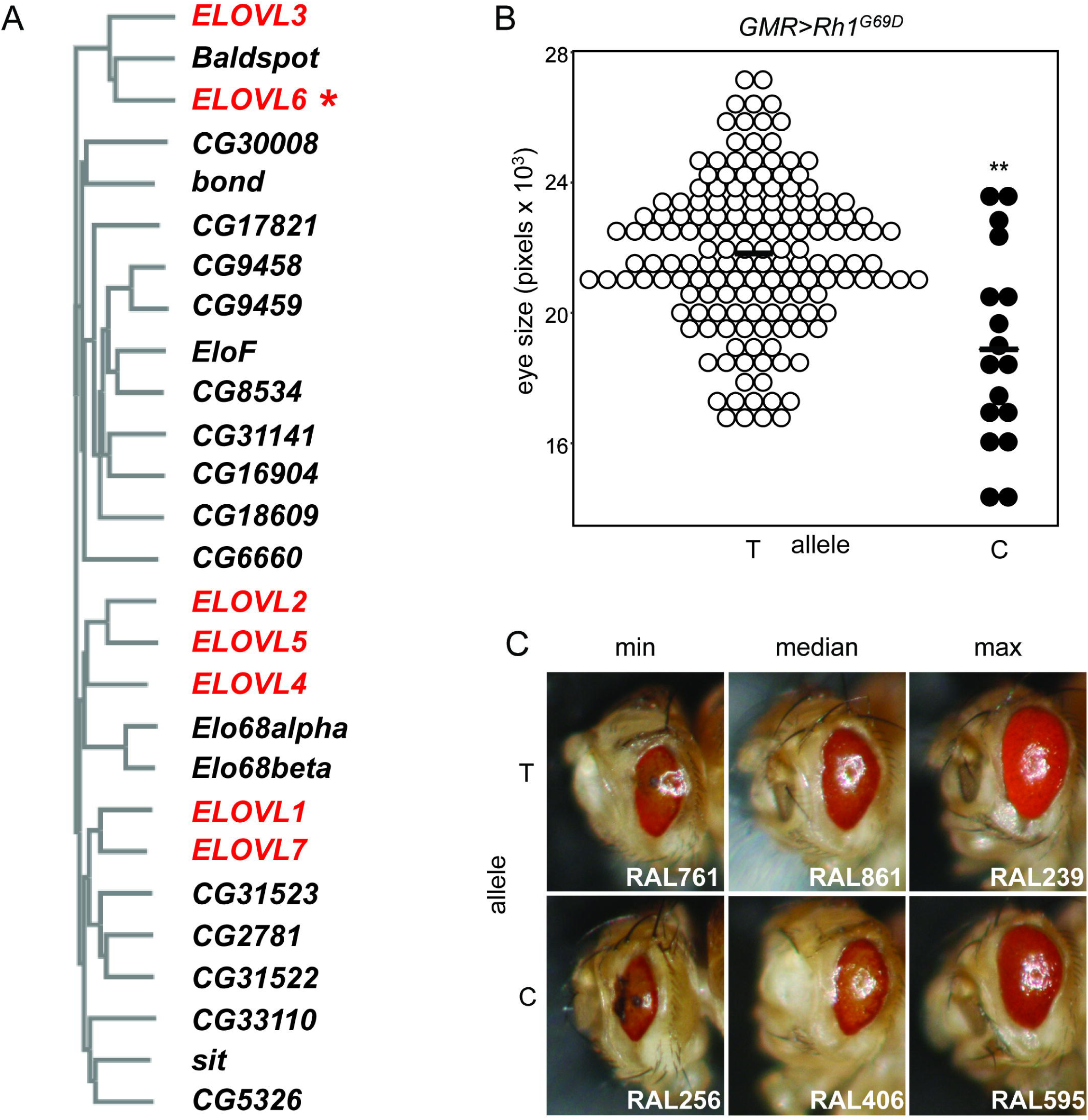
A SNP in *Baldspot* is associated with degeneration in a model of retinitis pigmentosa. **A.** *Baldspot* is the only *Drosophila* orthologue of mammalian *ELOVL6* (*). Phylogeny of long chain fatty acid elongases shows the relationship between *Drosophila* (black) and human (red) genes. **B.** The 3L:16644000 SNP in *Baldspot* (BDGP R5/dm3) is associated with *Rh1^G69D^*–induced degeneration in the DGRP. *Rh1^G69D^* DGRP eye size is plotted by allele identity. Strains carrying the minor C allele (18880 ± 3010 pixels) have significantly smaller eyes than those carrying the major, T allele (21822 ± 2294 pixels). **C.** Images from *Rh1^G69D^* DGRP strains that carry either the T or the C allele. Images were chosen from strains representing the range of eye sizes for the two alleles. All data from B and C were taken from Chow *et.al* 2016. ** P < 0.005

### Loss of *Baldspot* rescues *Rh1^G69D^*-induced retinal degeneration

To test whether eliminating *Baldspot* expression can modify *Rh1^G69D^-i*nduced retinal degeneration, we expressed an RNAi construct targeted against *Baldspot* in the *GMR-GAL4>UAS-Rh1^G69D^* background. Expression of this construct results in a strong, significant reduction in *Baldspot* mRNA (~7% of controls; P = 3.5 × 10^-3^) (Fig S2A). We found that loss of *Baldspot* expression results in partial rescue of Rh1^G69D^-induced retinal degeneration (*Rh1^G69D^/Baldspoti*) as compared to controls expressing only the Rh1^G69D^ misfolded protein and no RNAi (*Rh1^G69D^* controls) (Fig 2A). Quantification of eye size demonstrates a significantly larger, less degenerate eye in *Rh1^G69D^/Baldspoti* flies (13742 ± 913 pixels, n = 20) as compared to *Rh1^G69D^* controls (11432 ± 688 pixels, n = 20) (P = 5.3 × 10^-11^). We found a similar effect using another, independent RNAi strain (Fig S2B). Loss of *Baldspot* on a wild-type background (26478 ± 1191 pixels, n = 20) resulted in no significant change in eye size or phenotype as compared to genetically matched controls (26582 ± 1110 pixels n = 20) (Fig 2B). We conclude that loss of *Baldspot* expression provides significant rescue of *Rh1^G69D^-*induced retinal degeneration.

**Fig 2.**
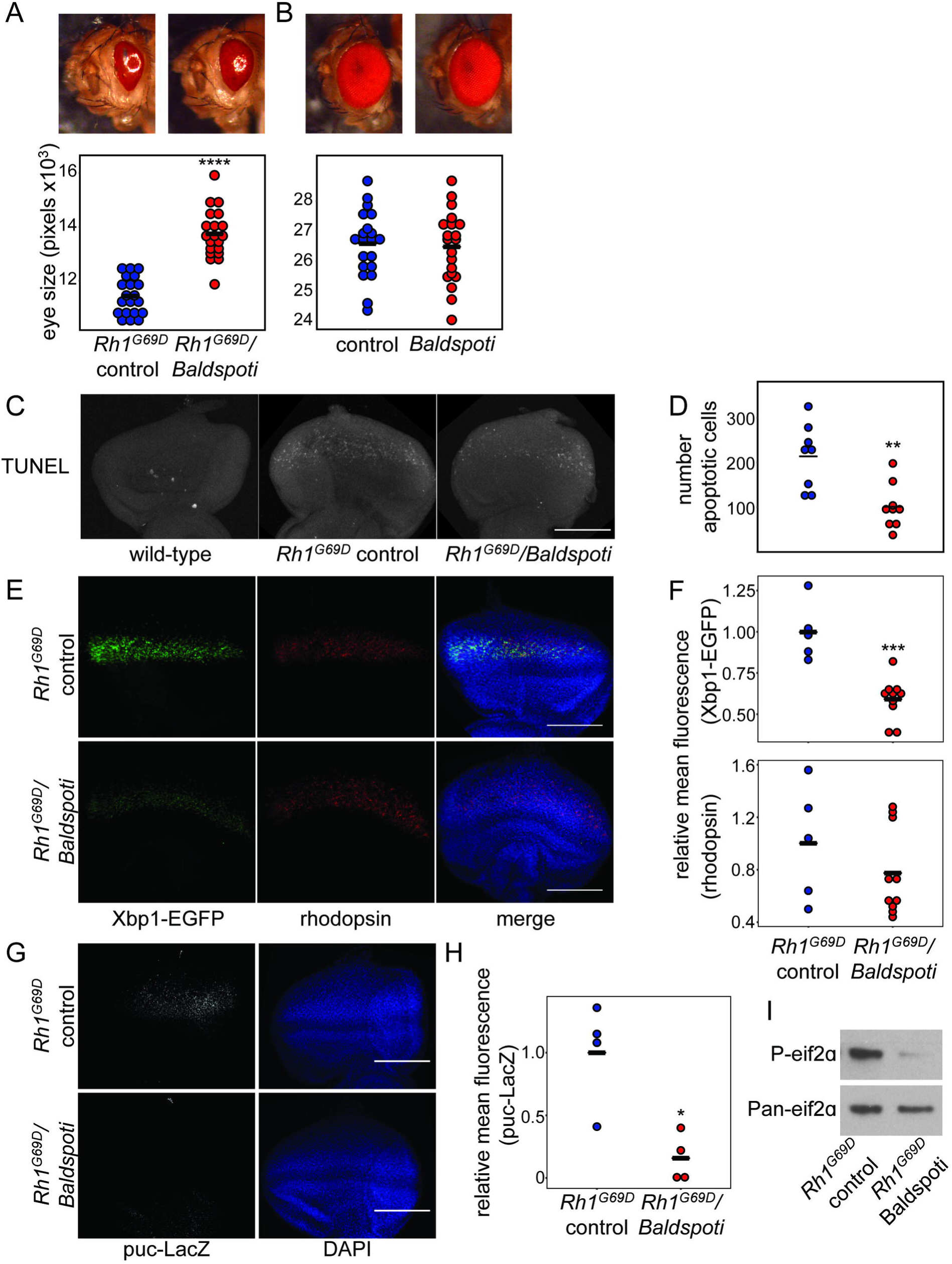
Loss of *Baldspot* in the *Rh1^G69D^* eye disc alleviates ER stress and apoptosis. **A.** Degeneration caused by overexpression of *Rh1^G69D^* is partially rescued by RNAi knockdown of *Baldspot* (*Rh1^G69D^/Baldspoti* vs *Rh1^G69D^* controls). This rescue was significant when eye size was quantified, as indicated by a larger eye in *Rh1^G69D^/Baldspoti* flies (14043 ± 960 pixels in *Rh1^G69D^/Baldspoti* flies and 11942 ± 473 pixels in controls). **B.** Loss of *Baldspot* on a wild-type background had no effect on eye phenotype (26478 ± 1191 pixels and 26582 ± 1110 pixels in controls). **C.** *Rh1^G69D^/Baldspoti* eye discs display reduced apoptosis levels compared to *Rh1^G69D^* controls, as measured by TUNEL staining. **D.** Cell death was quantified by counting the number of TUNEL positive cells in each eye disc. **E.** *Rh1^G69D^/Baldspoti* eye discs had reduced expression of Xbp1-EGFP as compared to *Rh1^G69D^* controls, while rhodopsin levels were unchanged. Eye discs were dissected from wandering L3 larvae expressing *Rh1^G69D^* and *UAS-Xbp1-EGFP*, stained for rhodopsin and GFP as an indicator of ER stress, and counterstained with 4’,6-diamidino-2-phenylindole (DAPI). **F.** Xbp1-EGFP levels were significantly reduced in *Rh1^G69D^/Baldspoti* eye discs (3.8 ± 0.8 fluorescent units) as compared to *Rh1^G69D^* controls (6.5 ± 1.1 fluorescent units). Rh1 levels were not changed (8.0 ± 3.5 fluorescent units vs. 10.4 ± 4.5 in controls). **G.** *Rh1^G69D^/Baldspoti* eye discs had reduced expression of puc-LacZ as compared to *Rh1^G69D^* controls. **H.** When quantified, LacZ levels were significantly reduced in *Rh1^G69D^/Baldspoti* eye discs (0.50 ± 0.6 fluorescent units) as compared to *Rh1^G69D^* controls (3.2 ± 1.3 fluorescent units). **I**. *Rh1^G69D^/Baldspoti* eye discs had reduced expression of P-eif2α as compared to *Rh1^G69D^* controls. Fluorescence was quantified using mean fluorescence from the image stack as calculated in Image J. Scale bars = 0.1 mm. * P < 0.05, ** P < 0.005, *** P < 0.0005, **** P < 0.0001.

### Loss of *Baldspot* in the *Rh1^G69D^* eye disc reduces ER stress and apoptosis

Rescue of eye size in the *Rh1^G69D^* model is often accompanied by reduction in apoptosis [29, 30]. Based on this and the rescue effect we observed, we hypothesized that apoptosis in the *Rh1^G69D^* eye discs would also be reduced upon loss of *Baldspot* expression. Indeed, we found that apoptosis, as measured by terminal deoxynucleotidyl transferase dUTP nick end labeling (TUNEL) staining, is significantly reduced in *Rh1^G69D^/Baldspoti* eye discs as compared to *Rh1^G69D^* controls (101 ± 95 TUNEL-positive cells in *Rh1^G69D^/Baldspoti* and 226 ± 69 TUNEL-positive cells in controls; P = 1.8 × 10^-3^) (Fig 2C-D).

Excessive ER stress in the *Rh1^G69D^* eye discs leads to apoptosis and degeneration and abnormal, degenerate adult eyes. To determine if the decrease in apoptosis we observed was due to changes in the ER stress response, we measured the activity of IRE1 and PERK, two of the transmembrane sensors of misfolded proteins in the ER. Under conditions that induce ER stress, *Xbp1* transcript is spliced by IRE1. The in-frame *Xbp1* transcript is translated and acts as a transcription factor, inducing genes involved in the UPR [8, 9]. The level of spliced *Xbp1* transcript is directly proportional to the degree of activation of the ER stress response. To measure IRE1 signaling and *Xbp1* splicing in the absence of *Baldspot* expression, we used a previously characterized transgenic *Xbp1-EGFP* marker [24, 31-33]. In this transgene, the 5’ end of the *Xbp1* mRNA is expressed under the control of the *UAS* promoter. The 3’ end of the gene, downstream of the IRE1 splice site, has been replaced with sequence encoding *EGFP*. As with the endogenous *Xbp1* transcript, the in-frame, spliced *Xbp1-EGFP* transcript, and subsequently the Xbp1-EGFP fusion protein, increases as ER stress signaling increases. Thus, EGFP signal increases as ER stress signaling increases. As previously reported, expression of *UAS-Xbp1-EGFP* in *Rh1^G69D^* controls results in increased EGFP, indicating high levels of ER stress (Fig 2E) [24, 31-33]. When *UAS-Xbp1-EGFP* is expressed in *Rh1^G69D^/Baldspoti* larvae eye discs, we see a significant reduction in EGFP signal (0.59 ± 0.13 in *Rh1^G69D^/Baldspoti* relative to controls, 1.00 ± 0.18; P = 9.6 × 10^-4^), indicating reduced ER stress signaling (Fig 2E-F). However, *Rh1^G69D^/Baldspoti* eye discs display no significant change in rhodopsin protein levels (0.78 ± 0.33 in *Rh1^G69D^/Baldspoti* relative to controls, 1.00 ± 0.44 in controls; P = 0.286) (Fig 2E-F), suggesting that loss of *Baldspot* expression reduces IRE1/Xbp1 signaling without affecting the accumulation or degradation of misfolded Rh1 protein.

IRE1-dependent cell death is primarily triggered by the Jun Kinase (JNK) signaling cascade when IRE1 has been chronically and strongly activated [7]. To measure activation of JNK signaling, we monitored expression of *puckered* (*puc*), a well characterized JNK target, using a LacZ-tagged allele (puc-LacZ). While puc-LacZ is induced in *Rh1^G69D^* controls (1.00 ± 0.41), it is significantly reduced in *Rh1^G69D^/Baldspoti* (0.50 ± 0.60 relative to controls; P = 9.5 × 10^-3^) (Fig 2G-H), suggesting that the observed reduction in cell death in the absence of *Baldspot* could be due to reduced signaling through the IRE1-JNK signaling axis, although JNK activation could additionally be achieved through a number of alternative pathways and influence other downstream processes besides apoptosis [34, 35].

We also examined the effect of *Baldspot* loss on the PERK branch of the UPR. When PERK is activated, it phosphorylates the translation initiation factor eIF2α, which decreases the translation of most mRNA transcripts with canonical translation initiation sequences [7]. We monitored P-eif2α levels by western blot in eye/brain-imaginal disc complexes isolated from *Rh1^G69D^* control and *Rh1^G69D^/Baldspoti* larvae. P-eif2α levels were substantially reduced in samples isolated from *Rh1^G69D^/Baldspoti* compared with *Rh1^G69D^* controls, indicating that PERK activity is reduced in the absence of *Baldspot* (Fig 2I). Because the ATF6 branch of the UPR has not been extensively studied in Drosophila, we are unable to draw any conclusions about the activity of this branch of the UPR.

Our data demonstrate that loss of *Baldspot* expression reduces IRE1 and PERK signaling misfolded protein levels. This is consistent with previous reports demonstrating that the cellular concentration of stearate is associated with increased activation of the ER stress response through direct activation of IRE1 and PERK, independently from misfolded proteins [36-38]. IRE1 and PERK both contain a conserved domain that is embedded in the ER membrane and can detect and respond to changes in membrane lipid composition, leading to increased activation of these sensors [39]. Together, this suggests that the modifying effect of Baldspot is linked to its fatty acid elongation activity.

### Stearate supplementation bypasses the requirement for *Baldspot*

Baldspot converts palmitate to stearate [26, 27] and the absence of Baldspot function should result in lower levels of stearate. We hypothesized that lower levels of stearate underlies the rescuing effect, reduced ER stress response, and retinal degeneration we observed, and that this signaling could be restored if stearate were supplemented during development. To test this, we raised *Rh1^G69D^/Baldspoti* and *Rh1^G69D^* control flies on media with or without 10% stearate and measured eye size in adults. This diet has no effect on eye size in wild-type individuals (Fig 3A). Stearate supplementation resulted in increased degeneration and reduced eye size in *Rh1^G69D^/Baldspoti* flies (14033 ± 1553 pixels) as compared to those raised on standard media with no stearate (16846 ± 1699 pixels; P = 1 × 10^-7^) (Fig 3B). This increase in degeneration in *Rh1^G69D^/Baldspoti* flies was accompanied by an increase in IRE1 activity and spliced *Xbp1*, as evidenced by an increase in Xbp1-EGFP expression in eye discs (1.26 ± 0.08 on stearate relative to standard media, 1.00 ± 0.10; P = 3.4 × 10^-3^) (Fig 3C-D). Stearate supplementation had no effect on eye size, retinal degeneration, or Xbp1-EGFP signal in *Rh1^G69D^* control flies (13478 ± 1197 pixels on stearate and 12873 ± 1006 pixels on standard media; P = 0.502; 0.83 ± 0. 15 on stearate relative to standard media, 1.00 ± 0.21; P = 0.171) (Fig 3 B-D), likely because of the high level of ER stress signaling already present. High stearate levels are sufficient to induce ER stress signaling and bypasses the need for Baldspot activity.

**Fig 3.**
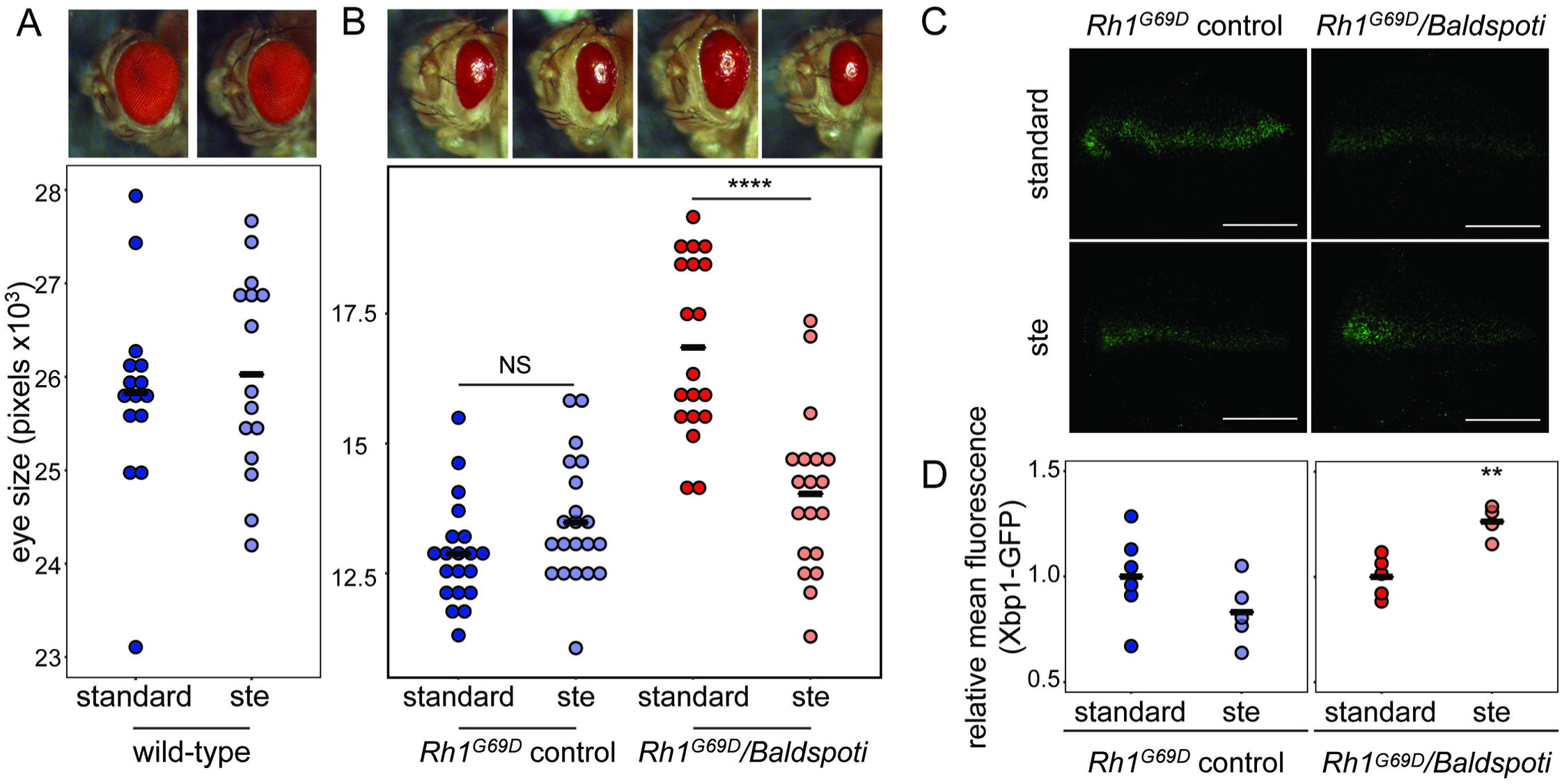
Stearate supplementation increases degeneration and activation of ER stress pathways in *Rh1^G69D^/Baldspoti* flies. **A.** Dietary supplementation with 10% stearate does not significantly impact eye size in wild-type flies (26026 ± 1089 pixels vs. standard media 25828 ± 1083 pixels). **B.** Eye size was significantly reduced in *Rh1^G69D^/Baldspoti* flies (*Baldspot* RNAi) when they were fed 10% stearate (14033 ± 1553 pixels) as compared to standard media (16846 ± 1699 pixels). Eye size was not significantly changed in *Rh1^G69D^* control flies (No RNAi) fed 10% stearate (13479 ± 1197 pixels) as compared to the standard media (12873 ± 1006 pixels). **C.** The 10% stearate diet also increased Xbp1-EGFP levels in eye discs from *Rh1^G69D^/Baldspoti* flies to a similar level seen in *Rh1^G69D^* controls. **D.** Xbp1-EGFP levels were significantly elevated in eye discs from *Rh1^G69D^/Baldspoti* flies (1.26 ± 0.08 relative to the standard media 1.00 ± 0.10). No change was observed in *Rh1^G69D^* controls (0.83± 0.15 relative to the standard media 1.00 ± 0.21). Scale bars = 0.1 mm. ** P < 0.005, **** P < 0.0001. NS = not significant.

### *Baldspot* is a general modifier of ER stress signaling

Because *Baldspot* is ubiquitously expressed (www.flyatlas2.org/), it is possible that *Baldspot’s* modifying role in the ER stress response is generalizable across different contexts and tissues. We first tested whether Baldspot modifies ER stress in another tissue. Because wing size is a visible and easily scorable trait, we used the *MS1096-GAL4* driver to express *Rh1^G69D^* in the developing larval wing disc (wing-*Rh1^G69D^*). The Rh1^G69D^ protein acts as a misfolded protein, inducing ER stress only in that tissue. The wing disc gives rise to the adult wing, and perturbations in cellular homeostasis in this tissue result in small, misshapen wings [40]. Expression of wing-*Rh1^G69D^* produces a phenotype similar to a vestigial wing (Fig 4A). When the RNAi construct targeted against *Baldspot* was concurrently expressed in the wing disc (wing-*Rh1^G69D^/Baldspoti*), this vestigial phenotype was partially rescued. The adult wings showed an increase in size and an increase in unfolding upon eclosion (Fig 4A). While this phenotype is variable in the degree of degeneration and unfolding of the wing, we do see a significant increase in 2D wing area and wing length in the absence of *Baldspot* (area: 100769 ± 9903 pixels in *wing-Rh1^G69D^/Baldspoti* and 57681 ± 30912 pixels in controls; P = 0.043; length: 633 ± 44 pixels in *wing-Rh1^G69D^/Baldspoti* and 378 ± 183 pixels in controls; P = 0.039) (Fig S3A-B). Similar to what we observed in the eye discs, this partial rescue was associated with a decrease in the level of Xbp1-EGFP in the wing disc (0.76 ± 0.17 in *wing-Rh1^G69D^/Baldspoti* relative to controls, 1.00 ± 0.14; P = 6.0 × 10^-3^) (Fig 4B-C). We also measured JNK activation in these wing discs by monitoring the *puc-LacZ* transgene. Again, as we saw in eye discs, LacZ expression is significantly reduced in wing discs lacking *Baldspot* (0.64 ± 0.12 in *wing-Rh1^G69D^/Baldspoti* relative to controls, 1.00 ± 0.12; P = 4.7 × 10^-3^) (Fig 4D-E). It appears that loss of *Baldspot* modifies UPR signaling and degeneration through the same mechanisms across different tissues and is not eye disc-specific.

**Fig 4.**
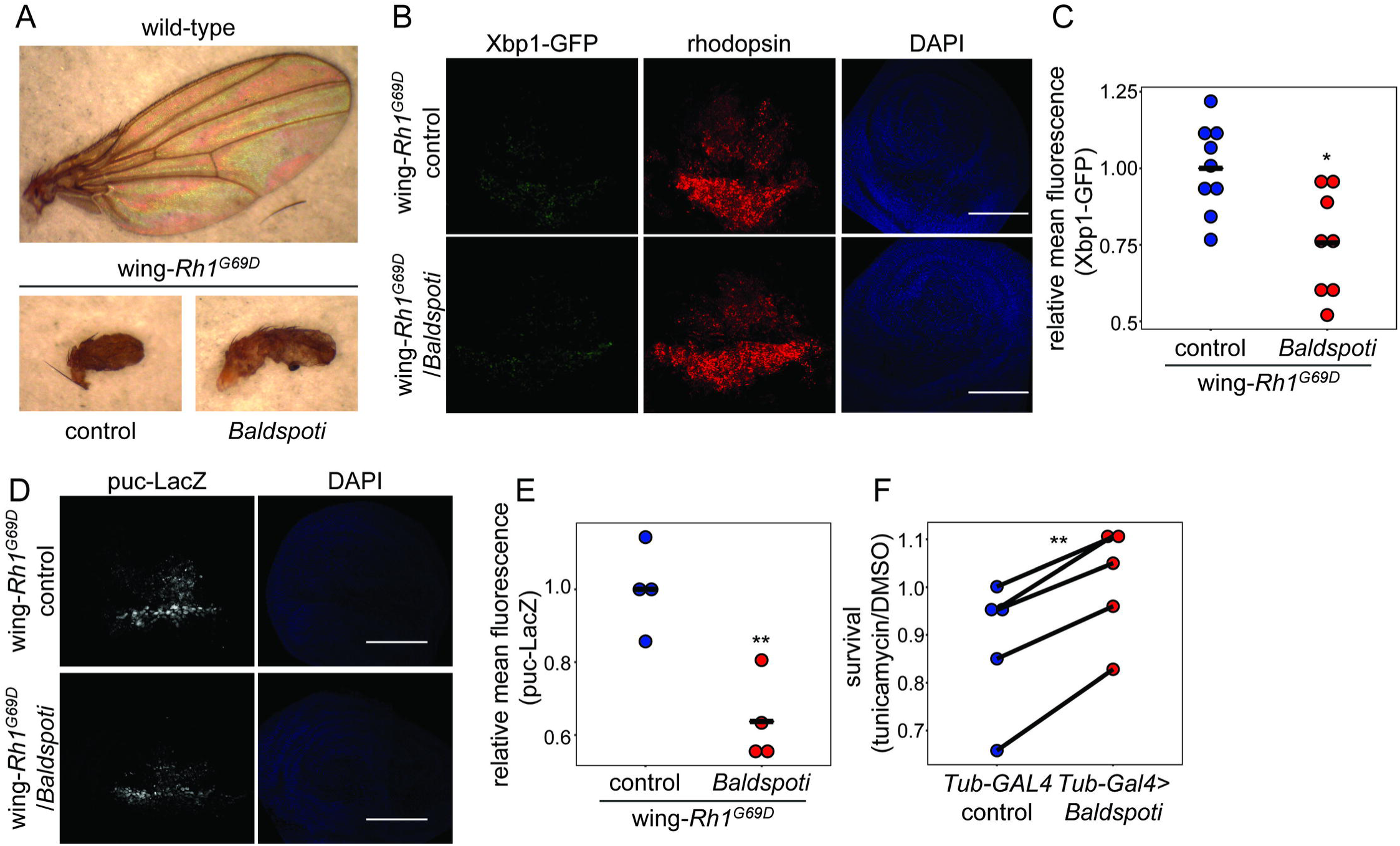
Baldspot is a general modifier of the ER stress response. **A.** Loss of *Baldspot* partially rescues the adult vestigial wing phenotype caused by expression of *Rh1^G69D^* in the wing disc (*wing-Rh1^G69D^/Baldspoti* vs wing-*Rh1^G69D^* controls). **B.** Wing discs from wing-*Rh1^G69D^/Baldspoti* flies had lower Xbp1-EGFP expression, as compared to wing discs from wing-Rh1^G69D^ controls. Wing discs were dissected from wandering L3 larvae expressing *Rh1^G69D^* and *UAS-Xbp1-EGFP* and stained for GFP and rhodopsin, with DAPI as a counterstain. **C.** Xbp1-EGFP levels are significantly reduced in *wing-Rh1^G69D^/Baldspoti* eye discs (0.76 ± 0.2 fold-change from controls) as compared to wing-*Rh1^G69D^* controls (1.00 ± 0.14). **D.** Wing discs from *wing-Rh1^G69D^/Baldspoti* flies had lower puc-LacZ expression, as compared to wing discs from wing-*Rh1^G69D^* controls. **E.** Puc-LacZ levels are significantly reduced in wing-*Rh1^G69D^/Baldspoti* wing discs (0.64 ± 0.12 fold-change from controls) as compared to wing-*Rh1^G69D^* controls (1.00 ± 0.12). **F.** Larvae with ubiquitous knockdown of *Baldspot* are significantly more resistant to Tunicamycin-induced ER stress than control larvae. L2 larvae were treated with tunicamycin or DMSO and survival to pupation was calculated as a measure of resistance to ER stress. Five representative experiments are shown. Scale bars = 0.1 mm. * P < 0.05, ** P < 0.005.

To test whether Baldspot modifies ER stress signaling by pharmacological induction, we used tunicamycin to induce ER stress. Tunicamycin inhibits N-linked glycosylation in the ER and results in massive misfolding, ER stress, and a robust UPR [41]. Treatment with tunicamycin in *Drosophila* larvae results in a robust activation of the ER stress response [33, 42]. We acutely exposed 2^nd^ instar control larvae (*Tub-GAL4*, no RNAi) and larvae with ubiquitous knockdown of *Baldspot* (*Tub-GAL4>Baldspot* RNAi) to tunicamycin or control DMSO and scored survival to pupation as a proxy for the physiological ability to recover from ER stress. To account for any larval lethality due to loss of *Baldspot*, we normalized the survival of larvae on tunicamycin to the DMSO treatment. Strikingly, larvae lacking *Baldspot* were less susceptible to tunicmycin-induced lethality than control larvae (Fig 4F, Table S1), as evidenced by increased survival to pupation (P = 9.2 × 10^-4^). This was accompanied by reduced *Xbp1* splicing (Fig S4A). We conclude that the modifying effect of *Baldspot* expression on the ER stress response is generalizable across different tissues and methods of ER stress induction.

### *Baldspot* alters RIDD activity of IRE1

Because our data indicates that Baldspot regulates Ire1/Xbp1 signaling and IRE1-dependent JNK signaling, we next tested whether Baldspot regulates another key IRE1 function: Regulated IRE1 Dependent Degradation of mRNAs (RIDD) [10, 11, 43]. In addition to *Xbp1* splicing, the RNase activity of activated IRE1 is also important for the degradation of a number of ER-associated transcripts through RIDD. To test if RIDD activity is affected by loss of *Baldspot* expression, we measured levels of known RIDD target mRNAs, under ER stress, in control cells and tissues with or without *Baldspot* expression. We employed S2 cell culture and treated S2 cells with dsRNA targeting either *Baldspot* or *EGFP* as a control. This treatment resulted in a near complete reduction in *Baldspot* transcript levels (~7.8% of control; P = 1.3 × 10^-9^) (Fig 5A).

**Fig 5.**
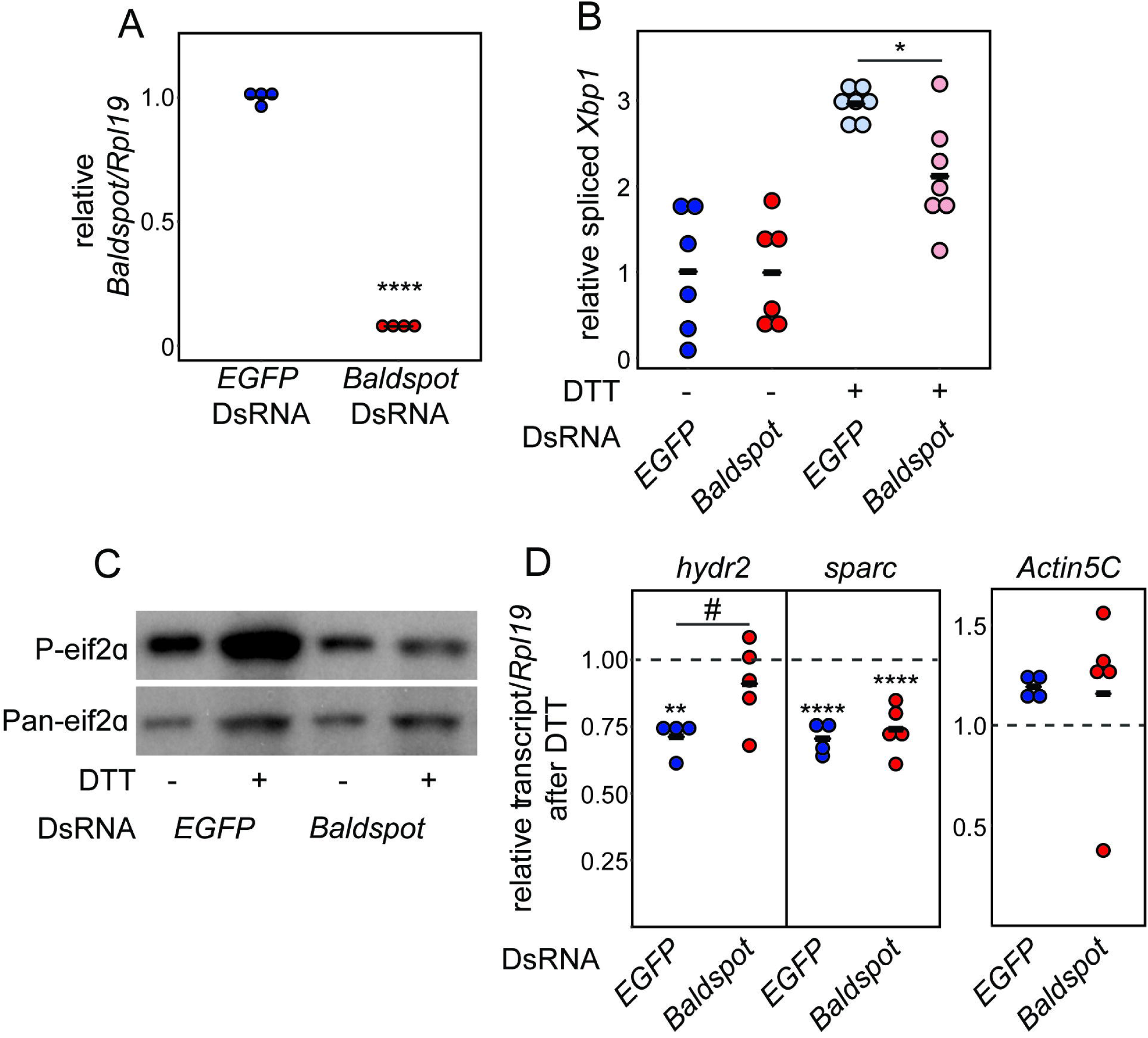
Loss of *Baldspot* reduces IRE1 RIDD activity. **A.** S2 Cells treated with DsRNA targeted against *Baldspot* had near complete reduction in *Baldspot* transcript levels (7.8%) as compared to control cells treated with DsRNA against *EGFP*. **B.** DTT treatment increased *Xbp1* splicing in control cells (2.96-fold increase), while splicing was reduced in cells treated with DsRNA against *Baldspot* (2.12-fold increase). **C.** DTT treatment increased phosphorylation of eif2α in control cells, but not in cells treated with DsRNA against *Baldspot*. **D.** *Hydr2* expression is significantly reduced in control cells treated with DTT (0.71 ± 0.07), but not in cells treated with *Baldspot* DsRNA (0.91 ± 0.16). *Sparc* expression is significantly reduced in both control and *Baldspot* knockdown cells treated with DTT (0.71 ± 0.06 in controls and 0.74 ± 0.09 in *Baldspot* knockdown). *Actin5c* transcript levels were unaffected by DTT treatment (1.19 ± 0.06 in controls and 1.16 ± 0.45 in *Baldspot* knockdown). The dotted line indicates transcript levels in cells treated with DMSO. * P < 0.05, ** P < 0.005, **** P < 0.0001, # P < 0.05.

We used Dithiothreitol (DTT) to induce ER stress in S2 cells. DTT is a reducing agent that breaks disulfide bonds in proteins, resulting in misfolded proteins and a strong ER stress response [44]. As expected, treatment of cells with DTT increased *Xbp1* splicing ~3-fold, indicative of ER stress. In agreement with our *in vivo* studies in the *Rh1^G69D^* fly, this splicing was significantly reduced in the absence of *Baldspot*, with only ~2-fold increase (P = 0.023), indicating that IRE1 signaling is disrupted (Fig 5B). We also monitored PERK activity by measuring levels of P-eif2α, which increase in control cells upon treatment with DTT (Fig 5C), as expected (P = 0.021; N = 3). In cells treated with *Baldspot* DsRNA, there was no increase in P-eif2α levels upon DTT exposure (P = 0.954; N = 3) (Fig 5C). Due to a lack of antibody, we could not monitor ATF6 activity in S2 cell culture. It appears that loss of *Baldspot* expression in S2 cells has a similar effect on ER stress response as observed in our *in vivo* model.

We next measured IRE1 RIDD activity in the absence of *Baldspot* in S2 cells. When RIDD is activated, ER-localized mRNAs are cleaved by the RNase domain of IRE1, leading to a reduction in protein translation into the ER [10, 43]. A number of these targets have been well-characterized in *Drosophila*, including the transcripts *sparc* and *hydr2*. Both of these mRNAs are degraded upon the initiation of ER stress [43, 45]. We tested the effect of *Baldspot* knockdown on these targets after treatment with DTT. *Hydr2* expression is significantly reduced in control cells treated with DTT (0.71 ± 0.07; P = 2.0 × 10^-3^), but not in those treated with DsRNA against *Baldspot* (0.91 ± 0.16; P = 0.41; P = 0.021) (Fig 5D). However, we found that *sparc* is equally degraded upon DTT treatment in cells treated with DsRNA against either *Baldspot* (0.74 ± 0.09; P = 2.0 × 10^-5^) or *EGFP* (0.71 ± 0.06; P = 1.9 × 10^-5^) (Fig 5D). Non-RIDD targets, such as *Actin5C*, were unaffected (Fig 5D). The fact that only some RIDD targets are affected is in agreement with our other results: loss of *Baldspot* activity partially reduces IRE1 functions, including RIDD.

## DISCUSSION

The ER stress response has the potential to be an important modifier of human disease. ER stress is activated in a wide variety of diseases ranging from Mendelian degenerative diseases to complex metabolic disorders [7]. The role of the ER stress response is complex and in some cases, a stronger response is beneficial and in other cases, a strong response underlies the disease. In some forms of retinitis pigmentosa, like the focus of this study, loss of vision is a direct result of the ER stress-induced cell death [46]. Neurodegenerative diseases such as Parkinson’s, Alzheimer’s, Amyotrophic Lateral Sclerosis (ALS), and prion disease are all associated with the accumulation of cytoplasmic misfolded proteins that can activate a secondary ER stress response and subsequent cell death [19]. In diabetes and obesity, misfolded proteins and the ER stress response are at least partially responsible for the cellular dysfunction that is a hallmark of the disease state [47]. In some cancers, however, cells selectively upregulate ER stress pathways associated with survival, while inhibiting those associated with cell death [48]. Genetic variation that alters the activation of the ER stress response may modify disease in the population.

We and others have shown that genetic variation can influence how and to what extent the ER stress response is activated. Genetic variation in *Drosophila* [22, 23], mouse [21], and human cells [20] can impact the expression of genes involved in ER stress pathways. Furthermore, this variation in expression has been linked to differences in the UPR and the downstream activation of cell death [21-23]. It is apparent that cryptic genetic variation drives these extensive inter-individual differences in the ER stress response [21]. Stress or disease conditions reveal the effect of this otherwise silent genetic variation. The same variation that drives differences in this important stress response might also influence disease heterogeneity. Indeed, modulating the ER stress response is an effective way of modulating disease outcome in models of human disease. [49-54].

In this study we show that loss of *Baldspot/ELOVL6* activity can affect the outcome of retinal degeneration and has the potential to be a broader regulator of ER stress-associated diseases. As a 16-to-18 carbon fatty acid elongase in the ER membrane, this enzyme is responsible for increasing the concentration of C18 stearate, a saturated long chain fatty acid [28, 37]. Increases in stearate concentrations in the ER membrane can induce and enhance ER stress, and is associated with disease outcomes in mammalian systems [36-38]. Here, we show that reducing the activity of Baldspot/ELOVL6 reduces the degree to which the UPR and apoptosis are induced in models of ER stress. Our study shows that Baldspot/ELOVL6 has a modifying effect on the IRE1 and PERK branches of the ER stress response. This is consistent with observations that increased long chain fatty acids in the ER membrane impacts IRE1 and PERK signaling [38, 39]. Future research will explore the Baldspot/ELOVL6 interaction with ER membrane composition and how this alters IRE1 and PERK oligomerization and function. It is particularly interesting to note that RIDD is only partially affected in the absence of *Baldspot*. In *Drosophila*, the RIDD activity of IRE1 targets most ER-localized mRNAs [43]. In contrast, mammalian RIDD is much more selective, and only mRNAs with specific signal sequences and RNA secondary structures are degraded by IRE1 [55]. More studies on how these processes are regulated in *Drosophila* and mammals is necessary to understand the differences in RIDD between species as well as the conservation of Baldspot/ELOVL6 function. These studies will enhance our understanding of how these pathways are regulated in different disease contexts.

Therapeutics that reduce *ELOVL6* activity could conceivably be used to treat or delay progression in a variety of ER stress-associated diseases. In RP patients with mutations that cause the accumulation of misfolded rhodopsin, similar to our *Drosophila* model, reduction of *ELOVL6* expression might reduce cell death, delaying vision loss. We present evidence that the modulation of the ER stress response through Baldspot/ELOVL6 activity may impact the severity of diseases beyond RP. Particularly intriguing is that *Baldspot/ELOVL6* can alter the ER stress response when it is induced by different conditions. Regardless of the mechanism by which ER stress is induced, inhibiting ELOVL6 activity reduces IRE1 and PERK signaling. In fact, in a recent study of the *db/db* mouse model of diabetes, loss of *ELOVL6* alters insulin sensitivity and the expression of ER stress genes [26]. It is likely that reducing *Baldspot/ELOVL6* expression would also alter complex diseases where ER stress contributes to degeneration and cell death. In some cases, even a small delay in degeneration could dramatically improve the quality of life.

We have identified *Baldspot/ELOVL6* in *Drosophila* as a novel modifier of the ER stress response. Its potential as a therapeutic target extends beyond RP to any number of ER stress-associated disorders. The identification of *Baldspot* in a natural genetic variation screen [23] suggests that variation in expression of this gene is well tolerated under healthy conditions and might be amenable to modulation. Identifying variable elements of the ER stress response pathway may prove to be a successful strategy to nominating new modifiers with broad applications to many ER stress-associated diseases.

## METHODS

### Fly stocks and maintenance

Flies were raised at room temperature on standard diet based on the Bloomington Stock Center standard medium with malt. The strain containing *GMR-GAL4* and *UAS-Rh1^G69D^* on the second chromosome (*GMR>Rh1^G69D^*) has been previously described [23]. The following strains are from the Bloomington Stock Center: *MS1096-GAL4* (8696), *Baldspot* RNAi (44101), control *attP40* (36304). The puc-LacZ enhancer trap is available from Kyoto (109029). A second *Baldspot* RNAi line (Vienna *Drosophila* Resource Center: 101557KK) showed similar rescue of the original *Rh1^G69D^* phenotype. Most analysis was performed in the Bloomington Stock Center RNAi line. The strains containing the *UAS-Xbp1-EGFP* transgenes were a gift from Don Ryoo (NYU) [24].

### Eye/wing imaging

For eye and wing images, adult females were collected under CO_2_ anesthesia and aged to 2-7 days, then flash frozen on dry ice. Eyes were imaged at 3X magnification using a Leica EC3 camera. Wings were dissected away from the body, then imaged at 4.5X magnification using the same camera. Eye area was measured in ImageJ as previously described [23].

### Immunohistochemistry

Eye discs and wing discs were dissected from wandering L3 larvae in cold 1X PBS, then immediately transferred to cold 4% PFA on ice. Tissues were fixed in 4% PFA for 15-20 min, then washed in 1XPAT (0.1% TritonX100) prior to blocking with 5% normal donkey serum. Tissues were stained with primary antibodies for rhodopsin (1:50, Developmental Studies Hybridoma Bank #4C5), GFP (1:2000, Thermo-Fisher #A6455), and LacZ (1:20, Developmental Studies Hydbridoma Bank #40-1a). Apoptosis was monitored using the ApopTag Red *In Situ* Apoptosis Detection Kit (Millipore #S7165). Tissues were mounted in Slowfade™ Diamond Antifade Mountant (ThermoFisher #S36967) and imaged with an Olympus FV1000 confocal microscope.

### Western blots

For DsRed blots, 10 brain-imaginal disc complexes were dissected from wandering L3 larvae and homogenized in 1X Laemmli/RIPA buffer containing 1X protease inhibitors (Roche cOmplete Mini EDTA-free protease inhibitor tablets). For GFP blots, 5 L2 larvae were homogenized in 1X Laemmli/RIPA buffer containing 1X protease inhibitors. For P-eif2α blots, protein from S2 cells was isolated in 1X Laemmli/RIPA buffer containing 1X protease inhibitors, as well as the phosphatase inhibitors Calyculin A and okadaic acid.

Equivalent amounts of protein were resolved by SDS-PAGE (10% acrylamide) and transferred to PVDF membrane by semi-dry transfer. Membranes were then treated with either 5% BSA or 5% milk protein block in 1XTBST prior to immunoblotting. Blots were probed with antibodies for RFP (1:500, Thermo-Fisher #R10367), GFP (1:5000, Thermo-Fisher #A6455), tubulin (1:2000, Developmental Studies Hybridoma Bank #12G10), P-eif2α (1:1000, abcam #32157), and Pan-eif2α (1:500, abcam #26197). Blots shown are representative of at least three biological replicates, and quantification was performed using Image J software.

### Stearate supplementation

Crosses were set up on the control, standard media alone, or standard media with 10% stearate supplementation. To make stearate supplemented media, standard media was melted in the microwave and kept at 98°C on a stir plate. Stearate (10% final concentration) was added to this media and mixed until lipids were homogenous with the media. The standard control diet received the same treatment, but no additional stearate was added. To determine retinal degeneration, flies were aged and imaged as described above. Antibody staining was performed on the eye imaginal discs of L3 larvae, also as described above.

### Tunicamycin treatment

Crosses to generate the indicated genotypes were set up on egg caps containing yeast paste. L2 larvae were then treated with either 10 μg/mL Tunicamycin (diluted 1:1000 from a 10 mg/mL stock solution) or 1:1000 DMSO in Schneider’s media for 30 minutes at room temperature. The larvae were then washed in 1XPBS twice and placed on solid media containing 6% yeast and 1% agar. Viability was determined by survival to pupation. Survival for each genotype was normalized to the DMSO-treated control condition. Each replicate represents ~25-60 larvae per genotype; a total 180 to 250 larvae per genotype were analyzed across all replicates. Protein was isolated from five L2 larvae expressing Xbp1-EGFP, one hour after treatment was concluded. A Western blot was performed to analyze Xbp1-EGFP levels.

### S2 cells

DsRNA was generated using the MEGAscript™ T7 Transcription kit (ThermoFisher #AM1334), with primers for *EGFP* (F: TTAATACGACTCACTATAGGGAGACCACAAGTTCAGCGTGTCC and R: TTAATACGACTCACTATAGGGAGAGGGGTGTTCTGCTGGTAGTG) and *Baldspot* (F: TTAATACGACTCACTATAGGGAGAATCCGCCCAGGTTCATCTCG and R: TTAATACGACTCACTATAGGGAGAGTCACATCTCGCAGCGCAAC). S2 cells were treated with DsRNA against *EGFP* (as a control) or against *Baldspot* at a density of approximately 2 × 10^6^ cells/mL in a 24-well plate. Cells were incubated with DsRNA for 4-7 days before being split and treated with either DMSO as a control or DTT. Concentrations of DTT varied with experiments: 0.5 mM for 1 hour (*Xbp1* splicing), 2.0 mM for 3 hours (RIDD target analysis), or 2.0 mM for 4 hours (P-eif2α detection). RNA was isolated from cells using either Trizol/chloroform extraction (*Xbp1* splicing) or the Direct-zol™ RNA Miniprep Kit (qPCR) and used to generate cDNA (Protoscript II, NEB). Protein was isolated from cells as described above and levels of P-eif2α normalized to Pan-eif2α as a loading control compared between matched DMSO or DTT-treated S2 cells.

Knockdown of *Baldspot* was confirmed using qPCR (primers: F: TGCTGGTCATCTTCGGTGGTC and R: ACGCAGACGGAGTGGAAGAG). *Xbp1* splicing was evaluated from the cDNA using PCR (primers for *Xbp1:* F: TCAGCCAATCCAACGCCAG and R: TGTTGTATACCCTGCGGCAG). The spliced and unspliced bands were separated on a 12% acrylamide gel and the proportion of these bands quantified using ImageJ software. RIDD target levels were analyzed by qPCR (*sparc* primers: F: AAAATGGGCTGTGTCCTAACC and R: TGCAGCACAATCTACTCAATCC; *hydr2* primers: F: CGCATACACGACTATTTAACGC and R: TTTGGTTT CT CTTTGATTTCCG; *Actin5C* primers: F: AT GT GT G ACG AAG AAGTT GCT and R: GAAGCACTTGCGGTGCACAAT; *rpl19* primers: F: AGGTCGGACTGCTTAGTGACC and R: CGCAAGCTTATCAAGGATGG). Transcript levels were normalized to *rpl19* and compared between matched DMSO or DTT-treated S2 cells.

### Statistics

Statistics were calculated using R software. P-values were determined using ANOVA for eye size, fluorescence levels, transcript levels in qPCR, and protein levels by western blot. Tukey multiple testing correction was used for the fatty acid feeding experiment and RIDD target analysis. Bonferroni correction was used for *Xbp1* splicing in S2 cells. A pairwise T-test was performed for larval tunicamycin treatment and P-eif2α response to DTT in S2 cells. A cutoff of P = 0.05 was used for significance.

## Acknowledgements

We thank Drs. Julie Hollien, Aylin Rodan, Anthea Letsou, and Carl Thummel for use of equipment and sharing reagents. We thank Emily Coelho for helpful comments.

---

**Supplemental Figure 1. The associated SNP in *Baldspot* is not significantly correlated with changes in expression. A.** Expression of *Baldspot* in strains carrying either the T or C allele of the 3L:16644000 SNP was determined from available RNA sequencing data in adult females [56]. *Baldspot* expression levels was not significantly impacted by this SNP. **B.** Expression of *Baldspot* was not significantly increased in brain-imaginal disc complexes isolated from DGRP lines expressing *Rh1^G69D^* and carrying the C allele, as compared to those carrying the T allele.

**Supplemental Figure 2. Confirmation of *Baldspot* RNAi. A.** The Bloomington *Drosophila* Stock Center *Baldspot* RNAi transgene (44101), used in most of this study, efficiently reduces expression of *Baldspot*. The RNAi construct was driven ubiquitously by *Tubulin-GAL4*, and expression determined in wandering L3 larvae. B. Degeneration caused by overexpression of *Rh1^G69D^* is partially rescued by knockdown of *Baldspot* using a second, independently derived RNAi line from Vienna *Drosophila* Resource Center (101557KK) (*Rh1^G69D^/Baldspoti* vs *Rh1^G69D^* controls). Similar to what was observed with the Bloomington *Drosophila* Stock Center strain, *Rh1^G69D^/Baldspoti* flies had larger, less degenerated eyes as compared to *Rh1^G69D^* controls (11323 ± 487 pixels vs. controls 10353 ± 798 pixels), indicating that the effects of *Baldspot* RNAi in this paper are due to loss of *Baldspot* expression. ** P < 0.005.

**Supplemental Figure 3. Wing size is significantly increased in *wing-Rh1^G69D^/Baldspoti* flies. A.** Loss of *Baldspot* significantly increases the size of wings expressing of *Rh1^G69D^* in the wing disc (100769 ± 9903 pixels vs. 57681 ± 30912 pixels in controls). **B.** Loss of *Baldspot* also significantly increases the length of wings expressing of *Rh1^G69D^* in the wing disc (633 ± 44 pixels vs. 378 ± 183 pixels in controls). * P < 0.05.

**Supplemental Figure 4. *Xbp1* splicing is impacted by tunicamycin treatment in L2 larvae. A.** Xbp1-EGFP protein levels are increased in *Tub-GAL4* control larvae one hour after treatment with tunicamycin as compared to DMSO-treated controls. Xbp1-EGFP is not increased in *Tub-GAL4/Baldspoti* larvae.

**Supplemental Table 1. Tunicamycin sensitivity.**

